# RprR is a plant-responsive regulator of exopolysaccharide production, biofilm formation, and virulence in *Ralstonia pseudosolanacearum*

**DOI:** 10.1101/2025.06.11.659228

**Authors:** Bridget O’Banion, Mariama D. Carter, Jose A. Sanchez-Gallego, Hanlei Li, Nicholas Wagner, Lan Thanh Chu, Loan Bui, Tuan Minh Tran, Caitilyn Allen

**Author notes:** These authors contributed equally to this work (Bridget O’Banion, Mariama D. Carter). Department of Biological Sciences, University of Missouri-Columbia, Columbia, Missouri, USA.

## Abstract

*Ralstonia pseudosolanacearum* (*Rp*s), which causes bacterial wilt disease of many crops, must integrate environmental signals to successfully transition from soil to its pathogenic niche in host plant xylem tissue. Mutating a putative sensing/signaling gene had little transcriptomic effect on *Rps* strain GMI1000 in culture. However, when the mutant grew in tomato over 180 genes were differentially expressed relative to wild type. The gene was therefore named *rprR* for *<u>R</u>alstonia* <u>p</u>lant-responsive regulator. *In planta*, the Δ*rprR* mutant dysregulated genes for diverse traits including stress response, degradation of phenolic compounds, motility, attachment, and production of extracellular polysaccharide (EPS), which is a key bacterial wilt virulence factor. Quantifying *Rps* EPS by ELISA found increased levels in stems of plants infected with Δ*rprR* as compared to wild type. Functional assays showed Δ*rprR* is defective in attachment to tomato roots, colonization of tomato stems, and bacterial wilt virulence. In rich medium, Δ*rprR* formed biofilm normally, but the mutant formed less biofilm in tomato stem homogenate and in tomato xylem sap under flow. This phenotype correlates with the mutant’s altered expression of EPS biosynthetic genes and aberrant extracellular matrix. When grown in tomato stem homogenate, Δ*rprR* produced 57% more of the bacterial signal cyclic di-GMP (c-di-GMP) than wild type. This is consistent with the presence in RprR of predicted c-di-GMP modulating domains. Together these findings reveal that RprR, which is highly conserved across plant pathogenic *Ralstonia*, modulates several bacterial wilt virulence traits in response to the plant host.

**Importance:** Members of the *Ralstonia solanacearum* species complex (RSSC) cause bacterial wilt, a globally destructive disease of market and subsistence crops. Like other plant-associated microbes, bacteria in the RSSC must integrate a complex array of biotic and abiotic signals to successfully infect plant hosts. RSSC genomes all encode an unusual protein, termed RprR, that contains multiple sensing and signaling domains, including two putative modulators of the secondary messenger c-di-GMP. Deleting RprR in *Ralstonia pseudosolanacearum* had a plant-dependent effect on many traits, including production of the key virulence factors biofilm and exopolysaccharide, as well as intracellular c-di-GMP levels. While c-di-GMP has been investigated in other plant pathogenic bacteria, this is the first report of its role in the RSSC. Most importantly, *rprR* was required for *Ralstonia* to effectively colonize plants and cause wilt disease. Thus, RprR is a plant-responsive sensor-regulator that controls pathogen adaptation to the host environment and virulence.

## Introduction

Bacteria use diverse sensors to integrate environmental stimuli into physiological responses that ensure fitness (1, 2). This is especially true for host-associated microbes that must adapt to dynamic feedback from the host. For example, members of the *Ralstonia solanacearum* species complex (RSSC), which cause bacterial wilt disease of many plants, have evolved intersecting regulatory networks to navigate a complex life cycle that includes survival in soil and water and explosive growth in the water-transporting xylem vessels of host plants (3, 4). In other well-studied bacteria, such networks are further linked with secondary messengers like bis- (3,5)-cyclic dimeric guanosine monophosphate (c-di-GMP), a cosmopolitan regulator of sessile-motile switches with further roles in plant and animal pathogenesis (5, 6).

The bacterial wilt disease cycle begins when a soil-borne RSSC cell senses and moves toward roots, then attaches and forms microcolonies on the root surface (7, 8). The pathogen then enters the root endosphere via wounds or natural openings, and migrates to the xylem tissue (7–10). Once inside xylem vessels, it must optimize its metabolism to this specialized nutritional environment, attach to vessel walls, form biofilms, and spread systemically throughout the host (11–14). RSSC members are well adapted to plant xylem, rapidly growing to densities greater than 10^8^ CFU/g stem. These large bacterial populations and their associated biofilm matrix fill xylem vessels and block water transport, leading to the characteristic wilting symptoms and death of the plant (15, 16). Swimming and twitching motility move RSSC cells to optimal sites; lectins and other adhesins facilitate attachment to plant surfaces; a consortium of extracellular enzymes degrade plant cell wall components and net-like plant DNA-histone traps; and dozens of Type III-secreted effectors suppress plant defenses and manipulate host metabolism to benefit the pathogen (4).

At several points in the bacterial wilt life cycle, bacteria in the RSSC develop and escape from biofilms. Biofilms, a common and successful lifeform, consist of microbial cells embedded in a self-produced extracellular matrix (ECM) that anchors cells, provides protection from a variety of stressors and improves nutrient uptake (17, 18). Plant pathogenic *Ralstonia* form biofilms during initial attachment and microcolony development on the root surface as well as during rapid growth following xylem invasion (7, 8, 12, 16). Members of the RSSC produce a distinctive extracellular polysaccharide called EPS I, which is a structurally complex heterogenous acidic polymer rich in N-acetylated sugars (19). EPS I, as well as extracellular DNA and proteinaceous lectins, are required for appropriate RSSC biofilm formation and are critical virulence factors (16, 20, 21). Accordingly, EPS production is tightly regulated and mutants that cannot make EPS are nearly avirulent (3, 21).

At each stage of the life cycle *Ralstonia* must fine-tune its gene expression to succeed. Inappropriately timed production of metabolic enzymes or virulence factors can waste resources and trigger host defenses. Despite decades of research on these systems, many elements of the *Ralstonia* regulatory network remain unknown (4). Among the best studied virulence regulators are HrpB and HrpG, which control production of the Type III secretion system (T3SS) and its effectors, along with hundreds of other genes (4). Another major regulator, the PhcA quorum sensing system, affects expression of 12% of genes in *R. pseudosolanacearum* (*Rps*) model strain GMI1000 and mediates a strategic shift in resource allocation between growth and virulence (11, 22). Interestingly, several *Ralstonia* regulators and virulence factors behave in a plant-specific manner. For example, PhcA represses HrpG expression at high cell densities in culture, but not when *Ralstonia* infects tomato stems (23, 24). Conditions present in the plant host also affect the activity of HrpG, PrhA, and the T3SS, but the specific signals and receptors remain elusive (4, 25, 26).

In addition to the dozens of known regulators, the genome of *Rps* GMI1000 encodes multiple putative sensors and regulators of unknown function. Among these is locus Rsp0254, which encodes a 129 kDa protein that includes six conserved domains predicted to function in signal perception and transduction, including cycling of c-di-GMP. A transmembrane HAMP domain suggests the protein’s location spans the periplasm and the cytoplasm. Previously, Rsp0254 was described as a light-responsive regulator of virulence based on the presence of a LOV (light, oxygen, voltage) sensing domain (27). An Rsp0524 deletion mutant was completely unable to colonize stems, which would make it avirulent. *In vitro* this mutant lacked any swimming and twitching motility and was defective in both EPS production and biofilm formation (27). These findings were unexpected for several reasons. First, this gene was not identified in several previous screens for avirulent mutants, although these screens repeatedly identified EPS biosynthesis genes, the virulence regulators described above, and the type II and type III secretion systems (21, 28–31). Further, there is reason to suspect that the described Rsp0524 mutant, which was not complemented, carries a second mutation in the *phcA* locus. The pleiotropic phenotype described by these authors closely resembles that of a *phcA* mutant (deficient in EPS and in virulence). The *phcA* locus frequently mutates spontaneously in culture (32–34). Finally, the conserved LOV domain was proposed to respond to blue light, a signal that is weak or absent in RSSC habitats like soil, roots, or stems.

To further define the role of Rsp0254, we independently constructed a mutant lacking this locus in *Rps* GMI1000. Transcriptomic and functional analyses revealed that although the protein has little importance *in vitro,* it has major effects when the pathogen is in the plant host. Rsp0254 was therefore named *rprR,* for *<u>R</u>alstonia* <u>p</u>lant-responsive regulator. While few genes were differentially expressed in Δ*rprR* under dark conditions or in blue light *in vitro*, this mutant dysregulated dozens of genes *in planta*. Many of the differentially expressed genes are involved in virulence, stress response, and EPS I synthesis. Consistent with this, *Rps* Δ*rprR* had altered EPS production, biofilm formation, and intracellular c-di-GMP levels, but only in plant-relevant conditions. Further, the Δ*rprR* mutant was defective in interactions with plants, exhibiting reduced root attachment, stem colonization, and virulence. Together these findings indicate that the highly conserved RprR protein regulates a wide array of functions required for *Rps* fitness during plant infection and pathogenesis.

## Results

### The *rprR* sequence and surrounding genomic region is highly conserved across the genus *Ralstonia*

To explore the conservation of *rprR* across the entire *Ralstonia* genus, we compiled 111 non-redundant genomes, including representatives from *R. pickettii*, *R. insidiosa*, *R. mannitolilytica*, and the four phylotypes in the *R. solanacearum* species complex (RSSC) (Table S1). Using as a query the *rprR* gene from RSSC model strain *Rps* GMI1000, we identified a putative homolog in 106 of the 111 genomes (>95% of genomes). A maximum likelihood tree of these *rprR* gene sequences aligns with the phylogeny of the genus *Ralstonia*, consistent with vertical inheritance from a common ancestral bacterium (Figure S1A).

The larger genomic context of *rprR* could provide clues to its biological function. Analysis of 12 representative *Ralstonia* genomes indicates that the synteny of this genomic region is also highly conserved (Figure S1B). All analyzed genomes contain genes neighboring *rprR* related to iron metabolism (*fur2*, *feoAB*, and a siderophore-interacting protein), energy/redox sensing (*aer2*), and reactive oxygen species (ROS) detoxification (*ahpCF*) (Figure 1A, Figure S1B, and Table S2). A smaller set of genes distinguish the plant-pathogenic RSSC from other *Ralstonia*. A c-di-GMP synthetase (*rsp0253*) is unique to the RSSC, while a three-gene cluster containing an alkylhydroperoxidase, an acetyltransferase, and an FMN-dependent NADH-azoreductase is present only in non-RSSC genomes (Table S2). Further, genes involved in hemin transport and degradation (*rsp0243-44*) were found only in RSSC phylotype I, IV, and a single phylotype II genome (Figure S1B).

**Figure 1:**
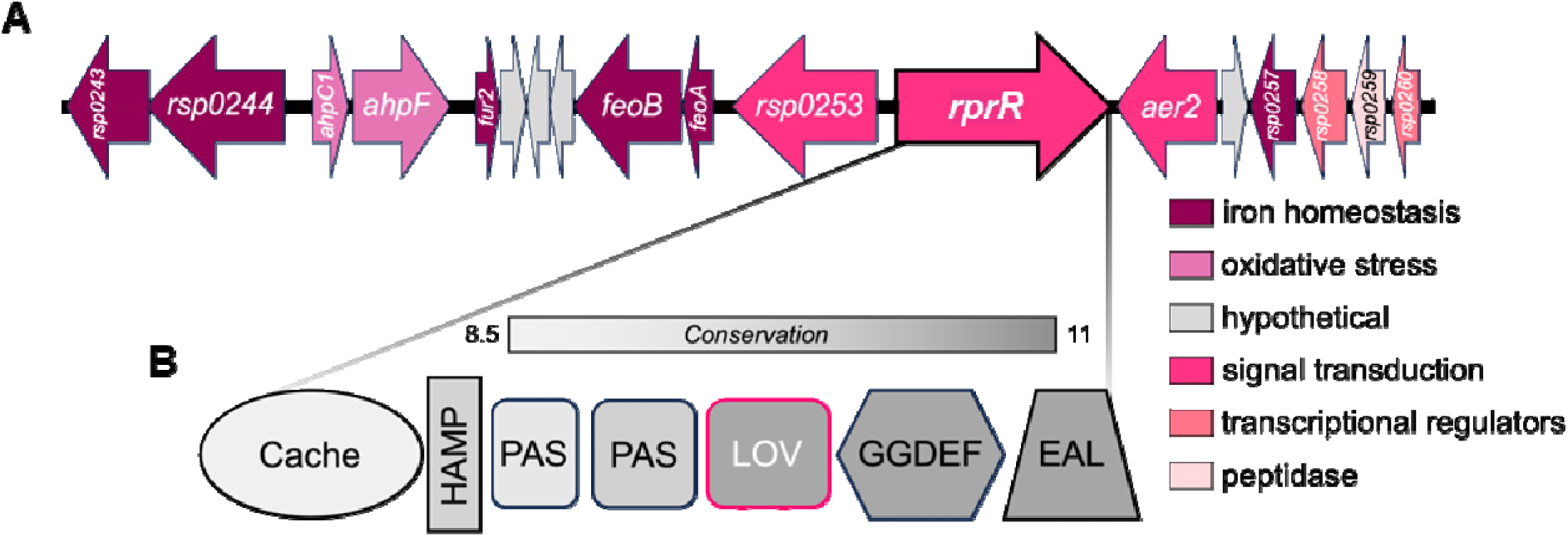
The multi-domain RprR protein and surrounding genes are conserved across the genus *Ralstonia*. **A)** Genes surrounding *rprR* (arrow with thicker border) on the *Rps* GMI1000 mega-plasmid. Arrow size represents approximate gene length. Gene arrows are color coded by functional category as indicated at right (see also Table S2). The *Rps* GMI1000 genome was manually visualized in the Joint Genome Institute Integrated Microbial Genomes web-server (35). **B)** The domain architecture of the predicted RprR protein. Domains are labeled and represented as different shapes. Shading corresponds to average amino acid conservation (Shenkin conservation score, calculated with default parameters in Jalview) within each domain boundary across 106 *Ralstonia* genomes (See also Table S1 and supplementary methods). The rounded rectangle labeled “LOV” denotes a PAS domain that encodes the characteristic LOV motif.

The *Ralstonia* RprR protein contains several sensory domains (Figure 1B). These include a periplasmic ligand-binding Cache domain and three cytoplasmic PAS superfamily domains, one of which contains the conserved LOV motif (36). A slight variant of the archetypal 8 amino acid LOV motif (GXNCRFLQ) is found in *Rps* GMI1000 *rprR* (GRNCRFLH) (37). RprR also contains two output domains related to the balance of c-di-GMP: EAL, which degrades c-di-GMP, and GGDEF, which synthesizes the signal (6). This domain architecture is unusual; many other bacterial LOV domain-containing proteins are histidine kinases and typically contain fewer additional signaling domains (38). We investigated the amino acid conservation in each of the RprR domains across all 106 *rprR*-encoding *Ralstonia* genomes. The EAL, GGDEF, and LOV domains were highly conserved, while the lowest level of conservation was observed in the Cache domain (Figure 1B).

### Blue light affects expression of very few *Rps* genes *in vitro*

We used RNA-seq to define the global effects of *rprR* and light in *Rps* GMI1000 and an in-frame *rprR* deletion mutant (hereafter Δ*rprR*). We compared transcriptomes of Δ*rprR* and wild-type GMI1000 *in vitro* in the presence and absence of blue light (Figure 2A). Strikingly, under blue light conditions only *rprR* itself was differentially expressed at q ≤0.05 between the genotypes. In dark conditions, two additional genes were also differentially expressed: a hypothetical gene predicted to encode for a small peptide (*rsc1470*) and a universal stress protein (USP, *rsp1561*) (Table S3). These three differentially expressed genes (DEGs) were all down-regulated in Δ*rprR*.

**Figure 2:**
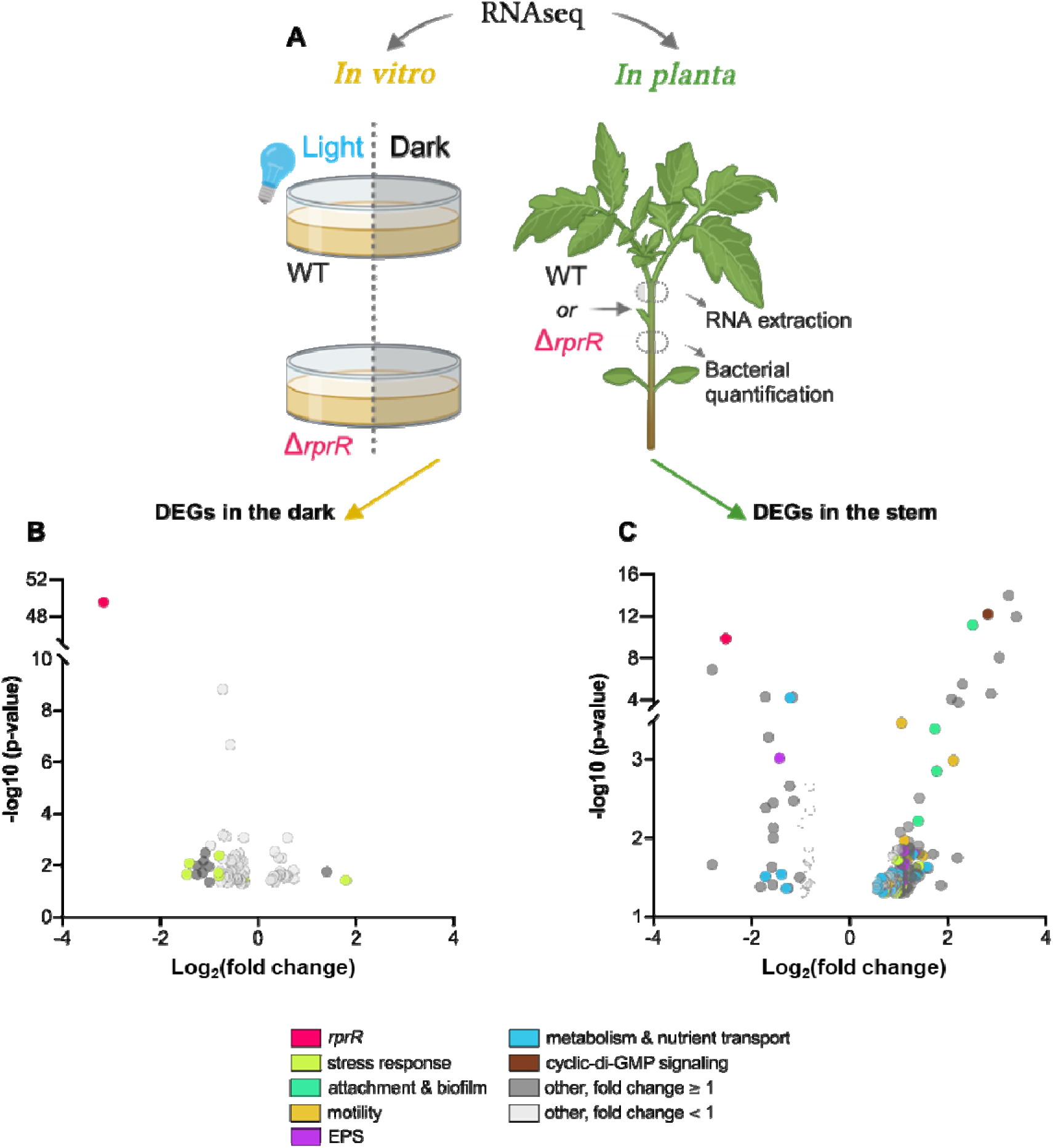
Mutating *rprR* dysregulates *Rps* stress response and virulence gene expression *in planta*. **A)** Design of experiments comparing gene expression of *Rps* GMI1000 wild-type (WT) and Δ*rprR* using RNA-seq. For *in vitro* (left), bacteria suspended in rich media were exposed to blue light or darkness for 2 hours (3 bio-replicates per treatment). For *in planta* (right), tomato plants were petiole-inoculated with each strain and 3 days later RNA was extracted from stem sections above the site of inoculation (4 bio-replicates per treatment). Stem samples below the site of inoculation were used to quantify bacterial colonization. **B-C)** Volcano plots showing *Rps* genes differentially expressed (DEGs, based on p-value) in Δ*rprR* compared to wild-type *in vitro* (B) and *in planta* (C). Panel B shows DEGs in dark conditions. Each circle represents a single gene and is color coded by functional categories shown in the legend. For genes that were not readily classified into listed categories, the corresponding circles are colored gray (light gray = log_2_ fold change < 1; dark gray = log_2_ fold change ≥ 1). Circles on the left of each graph represent genes with decreased expression in Δ*rprR*; those on the right are increased in Δ*rprR*. See also Tables S3 and S4.

For a larger view of potential transcriptional effects, we expanded our analysis to include DEGs with p ≤0.05. This larger set of genes included several involved in tolerance of reactive oxygen and nitrogen species (*oxyR* and *norB*) and other stress response mechanisms (melanin production and additional USPs) (Figure 2B, Table S3). Most of the less stringently defined DEGs, including those just listed, were differentially expressed between genotypes in the dark (90 total genes) rather than under blue light conditions (11 total genes). Further, only 85 genes were differentially expressed at p ≤0.05 in wild-type cells exposed to dark vs. blue light, and none of these met the more stringent q-value cutoff. Thus, blue light induces a minimal transcriptional change in cultured *Rps* and *rprR* may influence expression of *Rps* stress response genes.

### RprR influences *in planta* transcription of *Rps* genes for host-relevant behaviors

To assess the role of *rprR* in a more biologically relevant environment, we characterized *rprR*-dependent transcription *in planta* (Figure 2A). A larger set of *Rps* genes were differentially expressed in the Δ*rprR* mutant *in planta* than in culture, including genes for many virulence traits. At the stringent q ≤0.05 cutoff, expression of five genes decreased (coding for RprR itself, a probable lipoprotein, a putative transcriptional regulator, LdhA, and ExaC). Nine genes were upregulated in the Δ*rprR* mutant (coding for IbrAB, an EAL domain-containing protein, an N-acetyltransferase domain-containing protein, the adhesin RadA, a hypothetical transmembrane protein, a remnant transposase protein, a hypothetical TIR protein, and a putative transcriptional regulator). A p ≤0.05 cutoff identified 31 down-regulated and 156 up-regulated genes in Δ*rprR in planta* (Figure 2C, Table S4). The mutant increased expression of genes involved in general stress response (*rpoS*, USP [*rsp1561*]), degradation of phenolics (*fca*, *fcs*, *pcaJ*, *pcaG*), attachment (*radA*, *lecX*, *tadG2*), and motility (*fliC*, *flgE*, *pilE2*). Genes encoding probable lipoproteins (*rsc2609*, *rsc3141*) were down-regulated in Δ*rprR*, and c-di-GMP modulating enzymes (*rsc3143*, *rsp1623*) had variable expression patterns between the bacterial genotypes, with some increased and some decreased in Δ*rprR*. Notably, expression of the entire exopolysaccharide (EPS) I biosynthesis cluster (*epsABCDEFP*) increased in Δ*rprR*, consistent with the reduced expression of the negative EPS regulator *epsR*.

Together, these comparative genomic and transcriptomic analyses suggest *rprR* encodes a sensory protein that alters intracellular c-di-GMP levels in response to extracellular cues, thereby modulating *Rps* stress response and virulence. We next used functional assays to test specific hypotheses about the role of *rprR* in these behaviors. Because *in vitro* RNA-seq indicated that blue light does not significantly alter the *Rps* transcriptional landscape, light cues were not further studied.

### RprR influences EPS quantity, ECM quality, and cell fragility

We quantified the EPS produced by *Rps* cells growing *in vitro* and *in planta* to determine if expression of EPS biosynthetic genes correlated with biochemical measurements. An ELISA assay specific to a moiety in EPS I produced by members of the RSSC (16) detected no difference in EPS levels between wild-type and Δ*rprR* colonies grown on a rich medium (Figure 3A). In contrast, Δ*rprR* cells living in tomato stems produced significantly more EPS per cell compared to wild-type (Figure 3B). This plant-dependent change in EPS levels is consistent with the RNA-seq results. Complementing Δ*rprR* with the wild-type *rprR* gene under its native promoter (Δ*rprR+rprR*) partially restored the EPS production phenotype to wild-type levels (Figure 3B). A qRT-PCR analysis confirmed that *rprR* expression is restored to wild-type levels in the complemented mutant (Figure S2A), and expression of genes immediately surrounding *rprR* on the chromosome was not affected by the deletion (Table S3 and S4). Further, whole-genome sequencing of the wild-type, Δ*rprR*, and complemented Δ*rprR+rprR* strains revealed no additional mutations that could explain the observed phenotypes (Table S5).

**Figure 3:**
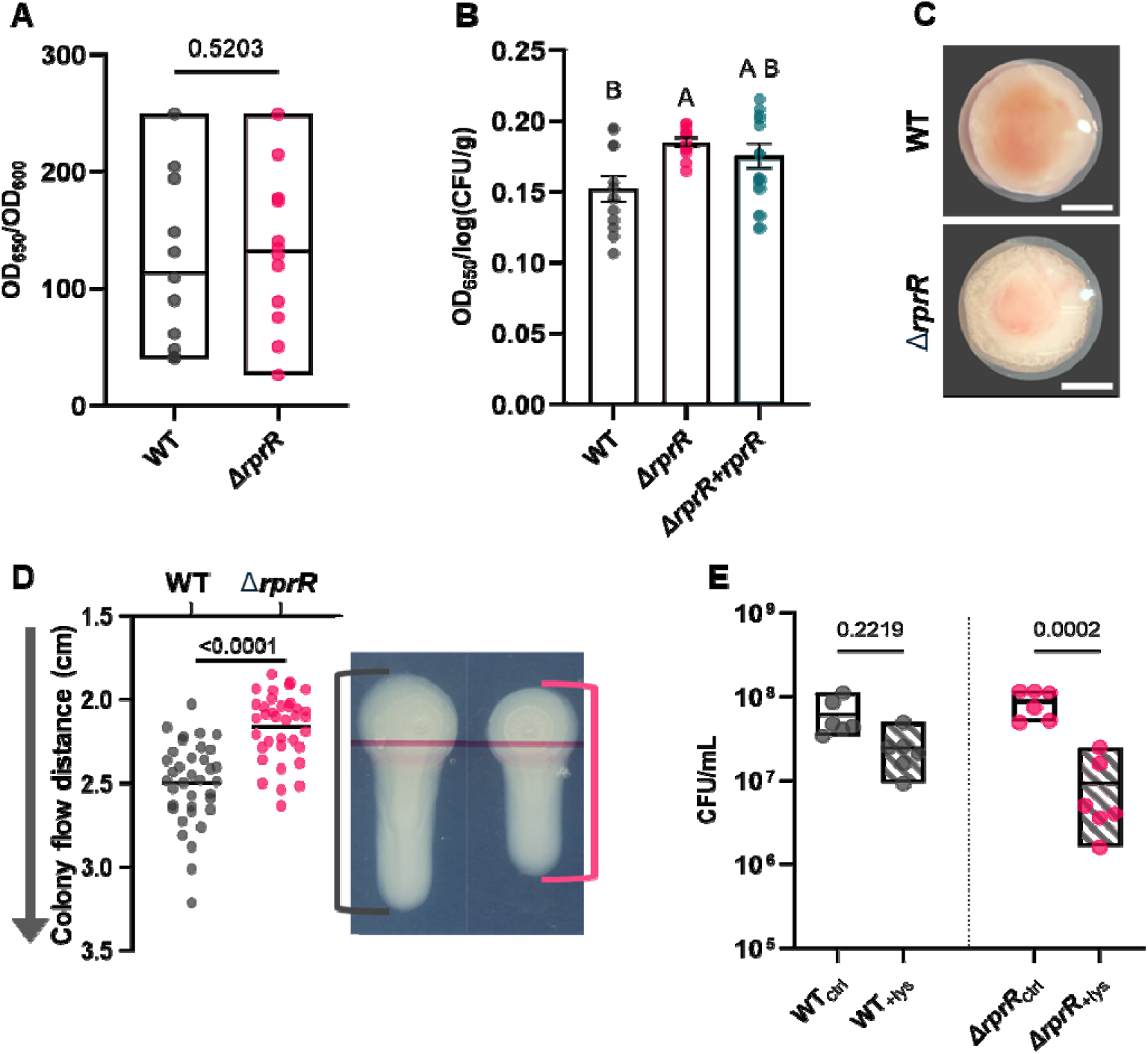
RprR influences EPS quantity, ECM quality, and cell fragility. Quantitative measurement of *Rps* exopolysaccharide production *in vitro* **(A)** *or in planta* **(B)**. Each circle represents the amount of an *Rps*-specific EPS I sugar moiety as determined by ELISA. **A)** *In vitro* EPS was measured in individual agar-grown colonies. Boxes show min to max values; horizontal lines indicate the mean. The experiment was repeated 3 times with 4 colonies each (n=12 replicates per treatment, indicated p-value by Welch’s t-test). **B)** Quantitative measurements of *Rps* EPS production *in planta*. Each circle represents EPS level in stem homogenate from a single infected plant stem. Bars show the mean and SEM. Different letter indicate significant differences between groups (p<0.05, Brown-Forsythe and Welch ANOVA).The experiment was repeated 3 times with 4 plants each (n=12 replicates per treatment). The differing Y-axis scales of panels A and B reflect distinct cell enumeration techniques (OD_600_ in colony suspension vs. log (CFU/g) for stem tissue). **C)** Colony morphologies of GMI1000 wild-type and Δ*rprR* on solid CPG media after three days at 28°C in the dark. Colonies were visualized at 10x magnification. The white scale bar indicates 2 mm. **D)** Gravitational flow measurements of *Rps* colonies. Each circle represents an individual agar-grown colony. Circles lower on the graph represent colonies with higher flow. Horizontal line indicate the mean. Representative images of wild-type and Δ*rprR* colonies after 60 seconds of vertical suspension are shown on the right. The horizontal pink line was to guide plate inoculation and does not represent a point of measurement. The experiment was repeated 3 times with 12 colonies each (n=36 replicates per treatment, p-value by Welch’s t-test). **E)** Population sizes of *Rps* strains following a 30 min exposure to 12.2 µg/mL lysozyme (+lys) or untreated (ctrl), as determined by serial dilution plating. Each circle represents the CFU/ml of a single microtiter well. Floating bars represent min to max values with line at the mean. Data reflect three independent experiments with two replicates each (n=6 per treatment, p-value by Mann-Whitney test).

Although Δ*rprR* and the wild-type strain produced similar EPS levels *in vitro* as determined by ELISA, we noticed that Δ*rprR* colonies on agar plates were qualitatively different from those of the wild-type parent. When grown on a solid rich medium, Δ*rprR* colonies were more translucent around the outer edges and had a curdled appearance (Figure 3C). RSSC bacteria produce a copious extracellular matrix (ECM) when grown on solid rich media. To test whether the ECM was physically different between wild-type and Δ*rprR,* we measured the flow rates of colonies when 3-day-old plates were tipped vertically. The Δ*rprR* colonies were significantly less fluid, suggesting the mutant’s ECM is qualitatively different from that of the wild-type strain (Figure 3D). Importantly, the cell density of colonies was equivalent between the two strains on a diluted version of the same media (Figure S2B). This suggests that the reduced fluidity of Δ*rprR* is due to a physical difference in the matrix, rather than a result of fewer matrix-producing bacteria.

The bacterial ECM can help protect against external stressors (17). We tested the hypothesis that the abnormal ECM of the Δ*rprR* mutant is associated with reduced physical resilience by exposing bacterial cells to the antibacterial enzyme lysozyme. Lysozyme, formally peptidoglycan *N*-acetylmuramoylhydrolase, degrades a primary structural component of bacterial cell walls, leading to cell lysis and death (39). We reasoned that a robust ECM normally protects *Rps* from the toxic effects of lysozyme, possibly by physically blocking the enzyme from reaching the bacterial cell wall or by supporting an enzyme-weakened cell wall until it can be repaired. Cells grown under the same conditions as for the tilt-plate assay described above were suspended in 12.2 µg/mL lysozyme for 30 minutes. This treatment caused only a non-significant population decrease in wild-type *Rps* cells, but it killed 89% of Δ*rprR* cells (Figure 3E). The lower survival following lysozyme exposure shows that Δ*rprR* cells are more fragile, possibly due to an abnormal ECM.

### *Rps* biofilm formation is modulated by RprR only in plant-relevant conditions

Biofilms, which are key to success at several points in the RSSC life cycle, are formed by cells embedded in an ECM (which is partially built of EPS) and are regulated by c-di-GMP in other microbes (17). Based on the c-di-GMP modulating domains of RprR and abnormal EPS levels and ECM fluidity of the Δ*rprR* mutant, we tested the hypothesis that *Rps* requires RprR for normal biofilm formation. Wild-type and Δ*rprR* cultures formed similar amounts of biofilm in a plate-based assay using rich medium, indicating that RprR does not play a role in biofilm formation under these conditions (Figure 4A). The contrasting *in vitro* and *in planta* RNA-seq and EPS quantification results led us to hypothesize that RprR specifically responds to host-derived signals to form biofilm. We tested this by incorporating filter-sterilized stem homogenate from healthy tomato plants into an *in vitro* biofilm assay. In the presence of this plant material, Δ*rprR* formed significantly less biofilm than wild-type (Figure 4B). This defect was fully restored in the Δ*rprR+rprR* complemented mutant strain (Figure 4B). An additional static biofilm assay using *ex vivo* xylem sap collected from healthy plants found that Δ*rprR* formed ∼26% less biofilm than wild-type *Rps* (Figure 3C). However, *Rps* cells live in actively transporting xylem vessels, which are not a static environment (12). We therefore used a microfluidic system that mimics xylem vessel surface chemistry, diameter, and flow rates to determine how flowing conditions would affect biofilm formation in tomato xylem sap (40). The addition of flow doubled the severity of the Δ*rprR* mutant’s biofilm defect, with these cells forming ∼58% less biofilm than the parental strain (Figure 3D-E). Together, these results indicate that although it has no effect in rich medium, RprR is required for normal *Rps* biofilm formation in conditions that chemically and physically mimic the plant host.

**Figure 4:**
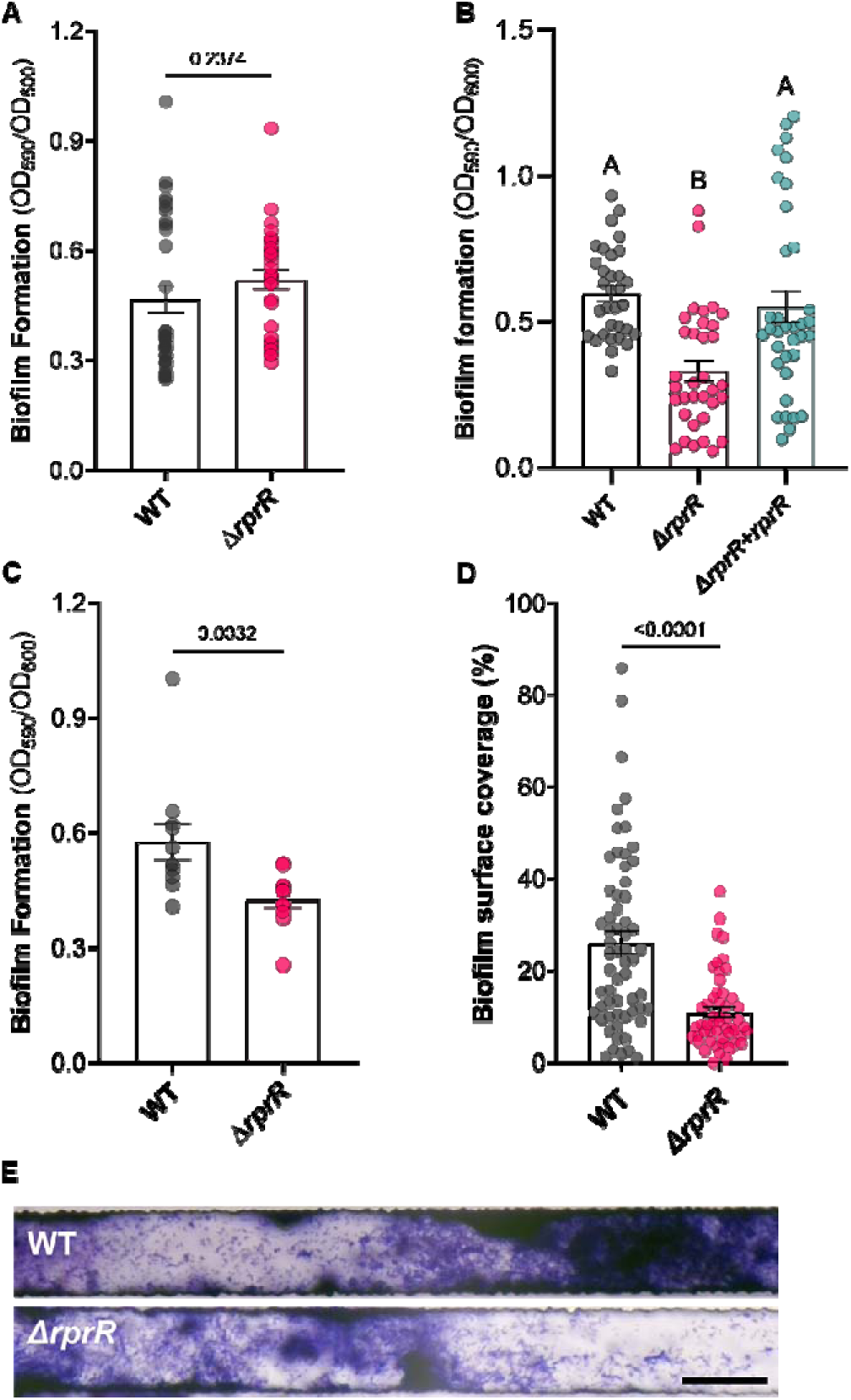
***Rps* biofilm formation is modulated by RprR only in plant-relevant conditions. A)** Biofilm formation in rich media as determined by a crystal violet-PVC plate assay. Each circle represents a single microtiter well. Bars represent the mean and SEM. Data shown reflect three independent experiments with 8-12 replicates each (n=32 per treatment, p-value by Welch’s T-test). **B)** Biofilm formation in filter-sterilized stem homogenate from healthy tomato plants. Each circle represents a single microtiter well. Bars represent the mean and SEM. Data reflect three independent experiments with 12 replicates each (n=36 per treatment). Outliers identified by a ROUT analysis (default parameters, implemented in GraphPad Prism) were removed prior to analysis and visualization. Different letters indicate differences between groups (p<0.05, Brown-Forsythe and Welch ANOVA). **C)** Biofilm formation in *ex vivo* tomato xylem sap. Each circle represents a single tube. Bars represent the mean and SEM. Data reflect four independent experiments with 2-3 replicates each (n=11 per treatment, p-value by Mann-Whitney test). **D-E)** Biofilm formation in *ex vivo* xylem sap under continuous flow. Each circle in **D** represents the crystal violet stain coverage % at a single location within a single channel (representative channel shown in **E**, with 50 µm black scale bar). The experiment was repeated twice, with 10 channels imaged for each device, at three different locations along each channel (n=60 per treatment, p-value by Mann-Whitney test). Outliers identified by a ROUT analysis (default parameters, implemented in GraphPad Prism) were removed prior to analysis and visualization.

### The Δ*rprR* mutant is impaired in host attachment, colonization, and virulence

Adhesion to the rhizoplane is the first physical interaction between plants and microbes and several adhesins were differentially expressed when the Δ*rprR* mutant grew *in planta* (41). We found that Δ*rprR* attached to tomato seedling roots ∼70% less frequently than wild-type *Rps* (Figure 5A). The mutant also had a significant defect in tomato stem colonization following direct inoculation via a cut petiole (Figure 5B, left). Tomato stems infected with Δ*rprR* contained ∼39% fewer *Rps* cells than those inoculated with the same number of wild-type cells. This defect cannot be explained by a metabolic defect in the xylem sap environment, as Δ*rprR* grows significantly better than wild-type in *ex vivo* sap (Figure S3A). Curiously, co-inoculating both strains into a single plant rescued the Δ*rprR* colonization defect (Figure 5B, right).

**Figure 5:**
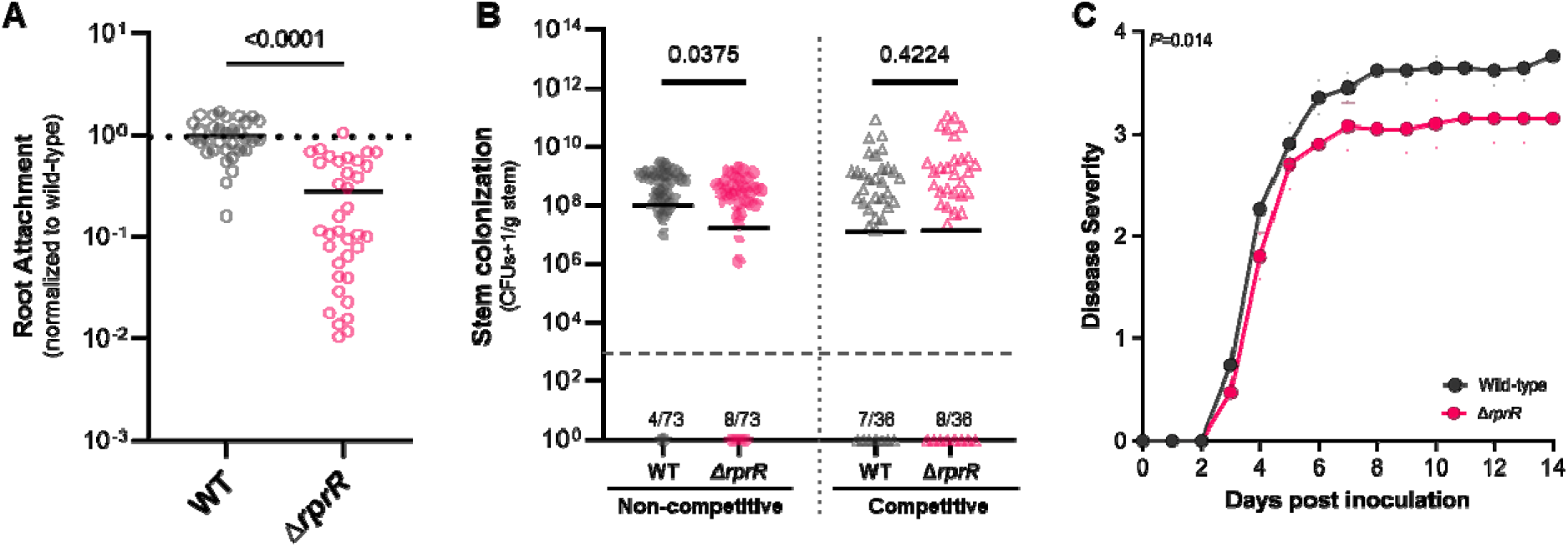
RprR is required for normal *Rps* root attachment, stem colonization, and virulence. **A)** *Rps* attachment to tomato seedling roots. Each symbol represents the population attached to 4-5 pooled roots, normalized to root biomass. Values were further normalized to wild-type levels; the black dashed line indicates value ‘1’ (mean wild-type attachment). Horizontal lines indicate mean attached population size. Data represent 4 independent experiments, each with 6-10 replicates (n=33-36 per treatment, p-value by Welch’s T-test). A single outlier identified by a ROUT analysis (default parameters, implemented in GraphPad Prism) was removed prior to analysis and visualization. **B)** *Rps* colonization of tomato stems under competitive and non-competitive conditions. Each circle represents the population in a single plant; lines indicate the geometric mean. The number of plants with undetectable *Rps* populations is indicated above the x-axis (limit of detection = ∼1000 CFU/g, represented by dashed line). A value of 1 was added to all final numbers to enable graphing ‘0’ values on a log scale. For non-competitive colonization (left), the two strains were individually inoculated into plant stems. The data represents four independent experiments with 15-23 plants each (n=73 per treatment, p-value by Mann-Whitney test). For competitive colonization (right), a 1:1 mixture of the two strains was inoculated into each plant stem. Data represent three independent experiments with 12 plants each (n=36 per treatment, p-value by Mann-Whitney test). **C)** Wilt disease progress on tomato following petiole inoculation, measured on a 0-4 disease index scale. Each symbol shows the mean disease index across three independent experiments, each containing 12-15 plants per treatment (n=40-42 plants per treatment, p-value by Repeated Measures two-way ANOVA). Error bars represent SEM.

The Δ*rprR* mutant’s altered gene expression, EPS production, biofilm formation, and host attachment and colonization behaviors suggested that RprR is required for full bacterial wilt virulence. We tested this hypothesis using two different virulence assays, which both showed that plants infected with Δ*rprR* had significantly decreased symptoms relative to wild-type *Rps* GMI1000. Following a naturalistic soil soak inoculation of unwounded tomato plants, wilt symptoms appeared more slowly on Δ*rprR*-inoculated plants than on plants inoculated with wild-type (Figure S3B). Similarly, when the root infection stage was bypassed by inoculating bacteria directly into xylem through a cut tomato leaf petiole, Δ*rprR* caused delayed wilting and also reached a lower final disease index than wild type (Figure 5C). In summary, four independent plant assays demonstrate that *Rps* needs the RprR protein to succeed at several different points in the disease process.

### Intracellular c-di-GMP levels are modulated by RprR in a plant-dependent manner

*Rps* RprR contains two domains predicted to modulate c-di-GMP, a cosmopolitan secondary messenger that regulates virulence in several pathogens (5). This molecule also often controls sessile-motile lifestyle switches and biofilm formation in bacteria (6). We observed an *rprR*-dependent biofilm defect when filter-sterilized healthy tomato stem homogenate was used as the growth substrate (Figure 4B). To explore the effect of plant defenses and wilt disease development on this behavior, we repeated the biofilm assay using stems from diseased plants infected with wild-type *Rps* or Δ*rprR*. The Δ*rprR* biofilm formation defect persisted under both of these diseased conditions, with the largest difference between wild-type and Δ*rprR* biofilm levels observed in stem homogenate from wild-type-infected tomato stems (Figure S4A-B). We hypothesized these plant-specific biofilm phenotypes were modulated by RprR-dependent changes in *Rps* c-di-GMP levels. To directly test this we quantified intracellular c-di-GMP levels in overnight cultures of wild-type and Δ*rprR* bacteria grown in filter-sterilized stem homogenate from wild-type-infected tomato plants, as described for the biofilm assays above. We also quantified c-di-GMP in overnight cultures grown in minimal medium. Supporting our hypothesis, Δ*rprR* produced 57% more c-di-GMP than wild-type when the bacteria were grown in stem homogenate (Figure 6). In contrast, deleting *rprR* had no effect on intracellular c-di-GMP levels in bacteria cultured in minimal media.

**Figure 6:**
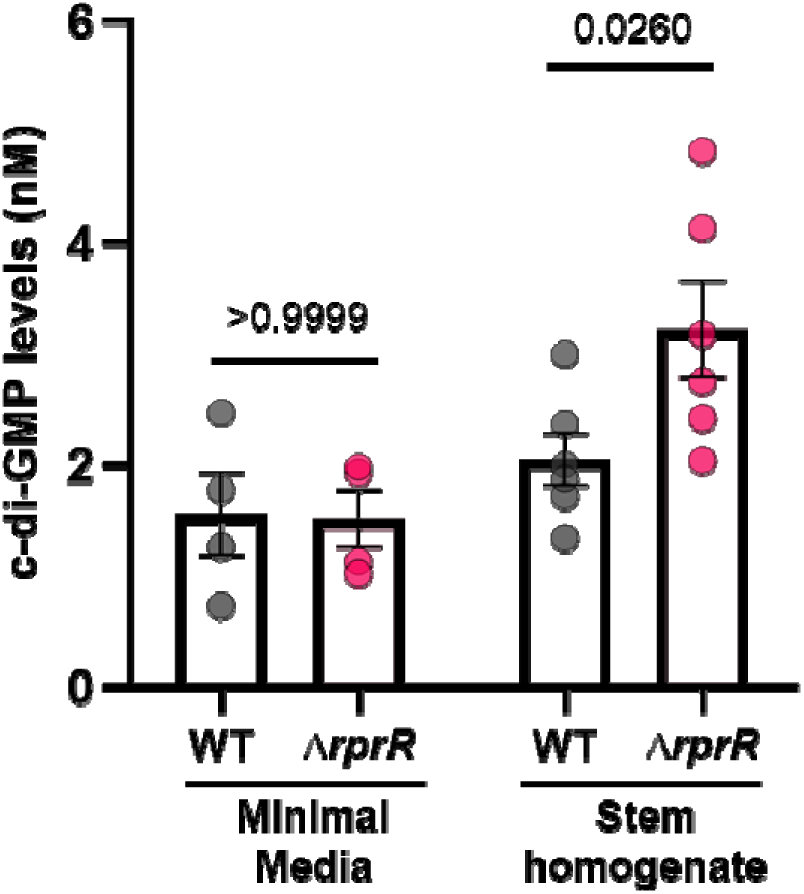
Deleting *rprR* increases c-di-GMP levels in response to plant extract. Concentration of c-di-GMP in *Rps* cells growing on different substrates. Each circle represents the nM concentration from within a single culture tube determined by LC-MS/MS. Bars represent the mean and SEM. For *Rps* grown in minimal media, the data reflect 4 independent overnight cultures for each strain (p-value by Mann-Whitney test). For *Rps* grown in filter-sterilized stem homogenate, the data reflect two replicate overnight cultures for three distinct batches of independently collected stem homogenate (n=6 for each strain, p-value by Mann-Whitney test).

## Discussion

The striking conservation of RprR across strains occupying diverse niches suggests this large multi-domain protein is important in *Ralstonia* biology. While LOV domain-containing proteins are typically studied as blue light photosensors with established roles in some foliar phytopathogens (42–45), *in vitro* transcriptomics revealed that blue light has only a minor effect on *Rps* transcription, and those effects are largely independent of *rprR*. Several assays conducted in the dark revealed an *rprR*-dependent phenotype, demonstrating a light-independent function for RprR in soil-borne plant pathogenic *Rps*. Because diurnal light cycles are inherently required for plant experiments, we cannot exclude an effect of light in our plant assays. Experiments using different light exposure methods could further define potential light-dependent and - independent roles for RprR in *Rps* biology.

Our transcriptomic and functional data show that RprR effects are highly context-dependent, making them difficult to capture via reductionist assays. Some *in vitro* studies directly contradicted results from whole plant assays. For example, some stress response genes were downregulated in the Δ*rprR* mutant *in vitro* but upregulated *in planta* (Figure 2B-C, yellow circles). It is not uncommon for host-associated microbes to behave differently outside their natural context (24), and mutating other *Rps* regulators has contradictory impacts on fitness *in planta* and *in vitro* (11, 46). We conclude that *in vitro* experiments are of limited value for understanding the true function of RprR. The plant-dependent role of *rprR* extends our knowledge of RSSC regulators critical for life *in planta*, which includes the previously characterized virulence and metabolic regulators PhcA and EfpR, and the plant-induced Prh and Hrp regulators of type III secretion (4, 11, 22, 46–54).

In addition to the LOV domain, the *Rps* RprR protein contains three non-photosensory signal input domains that likely incorporate multiplexed cues beyond light/oxidation. Additionally, LOV domain-containing proteins are known to exert light-independent transcriptional control (55). It is therefore not surprising that this work defined multiple light-independent roles for RprR. Mutating *rprR* had little effect *in vitro*, raising the possibility that the gene is not expressed in culture. However, previous transcriptomic studies found *rprR* is moderately expressed in rich medium with an absolute expression value of 9.74 and only slightly upregulated to 10.34 during growth in tomato plants; mutating *phcA* did not affect *rprR* expression *in planta* (11, 24). Further, our qRT-PCR results confirmed that the gene is transcribed *in vitro*. Although we cannot rule out the possibility that the RprR protein is not active *in vitro*, the presence of a periplasmic Cache receptor suggests that *rprR*-dependent regulation is triggered by a signal present in the plant but not in culture (56). Future studies investigating domain-specific effects on signal inputs and outputs could more specifically define the functional structure of RprR.

*Ralstonia* RprR contains two putative c-di-GMP active domains, suggesting it is involved in cycling this secondary messenger (6). The role of c-di-GMP has been investigated in several plant-pathogenic species (57), but for *Ralstonia* only the role of a different nucleotide secondary messenger (2’,3’-cGMP) has been reported (58). Still, the *Rps* GMI1000 genome contains nearly 40 genes predicted to modulate c-di-GMP, and biofilm formation plays a vital role in the pathogen’s life cycle (12, 57). In many bacterial species, high levels of intracellular c-di-GMP lead to increased biofilm formation, while lower levels correspond to a motile, planktonic state (6, 59). Biofilm formation is often driven via transcriptional, translational, or post-translational effects of c-di-GMP that increase production of specific matrix components such as polysaccharides and proteins (60–62). While *in vitro* transcriptomics detected no RprR-dependent effect on motility- or biofilm-associated gene expression, *in planta* RNA-seq analysis showed RprR modulates expression of genes involved in attachment and production of the biofilm matrix components EPS, adhesins, and lipoproteins. The effects of RprR on levels of EPS and intracellular c-di-GMP were similarly plant-dependent. These data indicate that plant pathogenic *Ralstonia* use RprR to modulate c-di-GMP signaling and ECM components in response to a plant-specific cue.

While cellular c-di-GMP levels and EPS production increased when *rprR* was deleted, the mutation decreased bacterial success at four critical steps of the *Rps* infection cycle: root attachment, biofilm formation, stem colonization, and virulence. Root attachment is dynamically dictated by extracellular polysaccharides, adhesins, pili, and other surface proteins (8, 63).

Expression of many genes related to these structures was altered in Δ*rprR*, including upregulation of the strong root attachment suppressor *lecX*, which could explain the observed adhesion defect (16). After breaking into host xylem vessels, *Rps* must successively produce and escape from biofilm to colonize, spread, and cause symptoms (20). Curiously, Δ*rprR*’s plant-dependent increase in c-di-GMP levels and EPS production did not translate to more robust biofilm formation. Rather, the mutant exhibited a significant biofilm defect specifically under plant-relevant conditions, and this defect was exacerbated under xylem-mimicking flow conditions. We suspect deleting *rprR* may cause structural differences in the ECM that impair proper biofilm formation. This is further supported by the mutant’s significant reduction in colony flow, stem colonization, and virulence. While Δ*rprR*-infected tomato stems contained over 1/3 less cells compared to wild-type-infected stems, co-inoculation with the parental strain rescued this defect. We suspect that co-inoculation complements the Δ*rprR* mutant by providing a common good in the extracellular space, specifically a properly functioning ECM.

The structural integrity of biofilms is influenced by the composition and scaffolding of the different matrix components (64–66). While we detected increased EPS production by Δ*rprR* cells, it is important to note that the ELISA detects a specific sugar moiety found in *Rps* EPS I. It does not provide information on correct assembly of EPS or other matrix components, which may also be altered in Δ*rprR*. For example, the Δ*rprR* mutant increases expression of a single adhesin *in planta*, while the genes encoding several lipoproteins and other membrane-associated proteins were downregulated. Similar proteins impact the structural integrity of biofilms produced by other bacterial species (67–69). Understanding how *Rps* builds its biofilm will require studies of the role of these proteins in *Rps* biofilm and their regulatory connection to RprR.

Deleting *rprR* reduced *Rps* stress tolerance. The Δ*rprR* mutant dysregulated various stress response genes and had a lower survival rate after acute lysozyme exposure. EPS and LOV domain-containing proteins influence bacterial stress response (17, 55, 70, 71), and RprR may link these behaviors in *Rps*. Bacterial EPS also modulates host recognition (72, 73). Curiously, wild-type GMI1000 formed significantly less biofilm in Δ*rprR*-infected stem homogenate than in stem homogenate from healthy, wild-type-, or Δ*rprR*+*rprR*-infected stem homogenate (Figure S4C). This suggests that infection by Δ*rprR* cells leads to a chemical environment that is less conducive to biofilm formation. The host plant may respond differently to Δ*rprR* cells, or the Δ*rprR* mutant metabolism may alter the host environment in ways that decrease biofilm potential. Comparative metabolomics of xylem sap from Δ*rprR-* and wild-type-infected plants could determine if RprR influences accumulation of antagonistic chemicals. Further, comparative host RNA-seq could reveal if the plant responds differentially to the mutant.

Adding a wild-type copy of *rprR* to the Δ*rprR* mutant at the chromosomal *att* locus restored wild-type levels of biofilm formation in stem homogenate and partially restored EPS production levels *in planta*. However, we could not complement several other phenotypes of Δ*rprR*. The complemented strain expresses *rprR* at wild-type levels and the genes surrounding *rprR* on the chromosome do not have altered expression due to the deletion. Further, whole-genome sequencing of the three strains revealed no additional mutations that could explain the lack of phenotypic complementation. Proper functioning of RprR may depend on its highly conserved genomic context. The *rprR* gene is flanked by the *aer* redox sensor and a GGDEF-encoding gene of unknown function, which both have regulatory inputs and outputs that could affect RprR (6, 74). We cannot rule out post-transcriptional or allosteric effects of protein-protein interactions that depend on appropriate subcellular localization of RprR (5). Further, spatially localized concentrations of c-di-GMP can affect phenotypes (75). If RprR affects a local pool of this molecule, this trait may not be restored by complementation at a remote genomic site.

In summary, this work establishes RprR as a critical sensor enabling *Rps* (and likely other plant pathogenic *Ralstonia)* to effectively cause bacterial wilt disease. Deleting RprR caused diverse phenotypes that reduced bacterial fitness in the dynamic host environment; these included dysregulation of various stress response and virulence genes, defective biofilm formation, and altered production of the critical wilt virulence factor EPS. Our data show that RprR acts across the disease cycle from early root attachment, through establishment in the stem, to end-stage disease outcomes. Notably, RprR is specifically plant-responsive. Further work is needed to identify the plant signal(s) that activate this regulator.

## Materials and Methods

### Bacterial strains and culture conditions

*Ralstonia pseudosolanacearum* (*Rps*) phyl. I-seq 18 strain GMI1000, Δ*rprR*, and Δ*rprR+rprR* were grown at 28°C in rich medium (CPG; 10 g/L peptone, 1 g/L casamino acids, 5 g/L glucose, 1 g/L yeast extract) or Boucher’s minimal medium (BMM; 3.4 g/L KH_2_PO_4_, 0.5 g/L (NH_4_)_2_SO_4_, 0.45 μM FeSO_4_ x 7H_2_0, 0.517 mM MgSO_4_, 10 mM glucose). *Escherichia coli* was grown at 37°C in LB (10 g/L tryptone, 5 g/L yeast extract, 10 g/L NaCl) with 25 mg/L kanamycin when appropriate. Unless otherwise indicated, *Rps* cultures were prepared as follows: Single colonies from a CPG plate were inoculated into CPG broth. Overnight cultures were centrifuged at 6-8,000 x g for 5-10 minutes to pellet cells. The supernatant was removed, and cells were resuspended in water. This wash step was repeated up to three times before measuring the optical density at 600 nm (OD_600_). This suspension was then used to inoculate various media to desired OD_600_ values.

### Plant maintenance and sap collection

Tomato plants (wilt-susceptible cv. ‘Bonny Best’) were grown at 28°C in a climate chamber with a 12-hour photoperiod as previously described (76). Xylem sap was collected from 5-week-old plants as described previously (76, 77), with minor modifications: after de-topping plants, sap was collected for 3 hours and pooled into a 50 mL tube on ice. Sap was filter-sterilized (0.22 µm), frozen at −20°C, and used within 3 weeks of collection.

### Transcriptomic analyses

*<u>in vitro</u>* <u>experimental design:</u> Three independent overnight cultures of wild-type GMI1000 and Δ*rprR* were grown in CPG to an OD_600_ 0.2-0.6. Washed cells were resuspended to a final OD_600_ of 0.2 in 8 mL of CPG and transferred to a small 60 mm petri dish to maximize even light exposure; dark treatment plates were wrapped in foil. Plates were shaken at 100 rpm under blue light (24-25 lux, wavelength range from 425-525nm with a peak of 464nm). After 2 hours, stop solution (5% phenol in ethanol) was added to the cultures and cells were centrifuged for 5 min at 10,000 rpm.

*<u>in planta</u>* <u>experimental design:</u> Wild-type GMI1000 and Δ*rprR* cultures were resuspended in water to a final OD_600_ of 0.001. Three-week-old plants were petiole inoculated with ∼2000 colony forming units (CFUs) of GMI1000 or Δ*rprR* (*76*). Three days post-inoculation, two ∼100 mg stem samples were collected from each plant: 1) a section above the site of inoculation was flash frozen in liquid nitrogen and stored at −80°C prior to RNA extraction, and 2) a section below the site of inoculation was used to quantify the bacterial population size (see below).

Samples with similar populations of wild-type (1.8-7.7×10^8^ CFU/g) and Δ*rprR* (1.5-7.6×10^8^ CFU/g) cells were selected to allow direct comparison of their transcriptomes. Selected samples were ground in stop solution using a PowerLyzer bead beater and centrifuged for 5 min at 10,000 rpm.

<u>RNA extraction, sequencing, and analysis:</u> Supernatant was discarded and total RNA was extracted from the pellet using the hot phenol-chloroform extraction method (24). RNA from each sample was extracted individually, measured for quality, and then 2 to 4 high quality samples were pooled per biological replicate. The *in vitro* experiment was repeated three times, and the *in planta* experiment was repeated four times. RNA samples (3 per *in vitro* treatment, 4 per *in planta* treatment) were sent to Novogene (Beijing) for library preparation, sequencing, and data analysis, as described previously (78). Briefly, libraries were generated using NEBNext Ultra RNA Library Prep Kit for Illumina (NEB, USA) and sequenced on an Illumina platform generating >8 million paired-end reads per sample. Reads were mapped to the *Rps* GMI1000 genome using Bowtie2 and DESeq2 was used for differential expression analysis. Due to incongruent gene expression compared to other replicates, one wild-type sample from the *in planta* dataset was removed from the final analysis. The gene expression datasets (Table S3 and S4) were manually filtered based on significance values (q-value<0.05; p-value<0.05) as indicated in the text. Data are available on NCBI in the Gene Expression Omnibus database (Accession Number GSE298975).

### Mutant construction

Plasmids, primers, and bacterial strains are listed in Table S6. Plasmids were extracted using a Zyppy Plasmid Miniprep Kit (Zymo). An unmarked in-frame deletion of *Rps* GMI1000 *rprR* was made using the *sacB* positive selection vector pUFR80 as described (79). Due to primer design constraints, 78 bp at the end of the 3,537 bp gene were not deleted. A deletion mutagenesis vector (pUFR80Δ*rprR*) was constructed with Gibson assembly and transformed into GMI1000 via electroporation (80). Gibson assembly was used to construct the complementation vector pRCK-*rprR*, which contains the wild-type *rprR* ORF and 186 bp of the upstream native promoter with flanking DNA to mediate insertion at the neutral chromosomal *att* site. This vector was transformed into Δ*rprR* via natural transformation, producing Δ*rprR+rprR*. The mutation and complementation were confirmed with PCR amplification and sequencing. For experimental assays using the complemented strain, a wild-type and Δ*rprR* strain containing empty pRCK inserts were used to control for effects of the vector itself. Additionally, Δ*rprR* was naturally transformed with an empty pRCG vector to create an additional antibiotic-resistant strain for competitive colonization experiments. All pRCK- and pRCG-containing strains were selected for using media + kanamycin or gentamycin at 25 mg/L as needed.

### EPS Quantification

*<u>in vitro</u>* <u>sample preparation:</u> *Rps* cultures were grown overnight in CPG broth. Turbid cultures were diluted 1:100 in sterile water and 5 µL drops were spotted on CPG agar plates and incubated in the dark at 28°C. After 3 days, a pipette was used to collect 10 µL of bacteria and their extracellular matrix from the middle of each colony into a microcentrifuge tube, vortexed with 1 mL sterile water, and diluted with sterile water to obtain an OD_600_ between 0.005-0.05.

*<u>in planta</u>* <u>sample preparation:</u> Plants were petiole-inoculated as described (76). Briefly, ∼2,000 cells of GM1000 or Δ*rprR* were applied to freshly cut leaf petioles of 21-day-old tomato plants. Four days post-infection, approximately 100 mg of stem directly above the cut petiole was homogenized in water using metal beads. A portion of the homogenate was immediately dilution plated to quantify CFU/gram, and the remaining homogenate was placed at −20°C until colonies could be counted. Samples with similar colonization levels between genotypes and across replicates (2.8×10^8^ − 3.9×10^9^ CFUs/g) were selected for ELISA quantification.

<u>ELISA:</u> EPS was measured immunologically with the AgDia DAS-ELISA PathoScreen^®^ Kit for *Ralstonia solanacearum* (PSP 33900, AgDia, Inc). This proprietary serological assay detects a conserved sugar moiety uniquely present in the exopolysaccharide of the *R. solanacearum* species complex. The manufacturer’s protocol was followed, with the following modification: a 10x concentration of the General Extract Buffer (GEB) was added at a 1:10 ratio to samples (10 µL 10x GEB + 90 µL sample) for a final 1x concentration. A negative water control was included, and the final OD_650_ readout for the negative control was subtracted from all experimental treatment wells prior to normalization by the OD_600_ (*in vitro* samples) or log(CFU/g) (*in planta* samples).

### Colony gravitational flow measurements

Turbid overnight CPG cultures of *Rps* were diluted 1:100 in sterile water and 5 µL drops were spotted on CPG plates and incubated in the dark at 28°C. After 3 days, plates were flipped vertically for 60 seconds to allow the colonies to flow down the plate. After 60 seconds, the plate was returned to a horizontal position and imaged. The distance from the top margin of each colony to the bottom border of the ‘drip’ was measured using ImageJ.

### Biofilm formation

Rich media: Static biofilm assays using CPG media in 96-well PVC plates were performed in the dark as described (16) via staining with 0.1% crystal violet.

<u>Static xylem sap:</u> For assays done in *ex vivo* xylem sap, strains were prepared as described above and resuspended to an OD_600_ 0.01 in xylem sap. Suspensions were aliquoted (1.4 mL/tube) into 4mm borosilicate glass tubes. The top of each tube was wrapped in parafilm and a loose-fitting cap was placed on top. Tubes were incubated statically at 28°C in the dark. After 5 days, the suspension was gently mixed using a pipette and the OD_600_ was measured.

Biofilm staining with 0.1% crystal violet was performed as described (16), except that tubes were washed only once with water prior to de-staining.

<u>Stem homogenate:</u> Homogenized and filter-sterilized stem from healthy and *Rps*-infected tomato plants was used as a growth medium for biofilm assays. Three days after 21-day-old plants were petiole inoculated (76), approximately 200 mg of stem directly surrounding the cut petiole was collected from each plant and homogenized in water using metal beads. For healthy plants, stem tissue was collected from 24-day-old plants to control for age. Homogenized samples for each treatment were combined (n=6-13 plants per batch) and sterilized with a 0.22 µm filter. Stem samples for all treatments were collected in three separate biological replicates, for three batches of pooled stem homogenate. The ratio of plant material to water was maintained across treatments and replicates to control for the density of plant substance within each batch.

For stem homogenate biofilm assays, *Rps* cultures grown overnight in CPG broth were diluted 1:100 in sterile water. Thawed stem homogenate was added to a 96-well plate (140 µL/well; Celltreat #229590) and 10 µL of the diluted cell suspensions were added. The plate and lid were wrapped in parafilm and incubated at 28°C in the dark. After 3 days, each well was gently mixed using a pipette and the OD_600_ was measured. The wells were washed, stained, and analyzed as for ‘static xylem sap’ above.

<u>Xylem sap under flow:</u> Microfluidic device chips were pre-coated overnight at 37°C with 10 µg/ml CMC-DOPA as described (16, 40). Overnight *Rps* CPG cultures were adjusted to 10^9^ CFU/mL with sterile water, then seeded into microfluidic devices by a syringe pump for 6 hours under static conditions. After seeding, unattached cells were flushed from the microfluidic system with sterile water. Filter-sterilized xylem sap from healthy plants was then pumped through the channels at 40 µL/hour for 72 hours in the dark. The sap-containing syringes only contained enough xylem sap for 24 hours, so fresh xylem sap was introduced each day. After 72 hours, the microfluidic devices were stained with 1% crystal violet and washed three times.

Images were captured using a compound microscope and analyzed with Fiji software to quantify biofilm surface coverage in the channels. Experiments were repeated twice, with 10 channels imaged for each device, at three different locations along each channel.

### Root attachment

Rhizoplane bacterial populations were measured as described (8, 76) with minor modifications: seedlings were flood-inoculated with 15 mL of 10^6^ CFU/mL *Rps* GMI1000 or Δ*rprR* suspended in water. The cell suspension was removed after 2 hours in the dark.

Seedlings were then flooded with sterile water, swirled for 10 seconds, and water was removed. Seedlings were cut to separate the root from the hypocotyl, blotted dry, and pooled (4-5 roots per technical replicate). Each pooled sample was weighed before homogenization and dilution plating to quantify attached cells.

### Virulence and stem colonization

Plant assays were performed as described (76). Briefly, 21-day-old plants were either soil soak-inoculated or petiole-inoculated with GM1000 or Δ*rprR* cells to quantify virulence. Symptom development was rated for 14 days on a 0-4 scale. For non-competitive stem colonization, plants were petiole-inoculated with ∼2,000 cells of GM1000 or Δ*rprR*. Three days post-infection, approximately 100 mg of stem directly below the cut petiole was homogenized in water, diluted, and plated to quantify CFU/gram stem. For competitive stem colonization, plants were petiole-inoculated with a 1:1 suspension of wild-type GMI1000-Km and Δ*rprR*-Gm. Stems were sampled similarly to non-competitive assays except that samples were serially dilution plated on both CPG + kanamycin and CPG + gentamycin to quantify CFU/g stem of each strain.

### Intracellular c-di-GMP measurements

Overnight bacterial cultures were resuspended in water at OD_600_ 0.001. A 100 µL aliquot of the GMI1000 or Δ*rprR* suspensions were used to seed culture tubes containing either 3 mL of BMM + 0.1% tryptone or 2 mL of filter-sterilized stem homogenate collected from tomato plants infected with wild-type GMI1000. The stem homogenate batches used for this experiment were the same as those used in the biofilm experiment. Because the stem homogenate came from infected plants, there was likely a baseline level of c-di-GMP in the mixture prior to inoculation. However, the baseline level was equivalent across treatments, so any differences observed were due to the behavior of the inoculated wild-type or Δ*rprR* cells. Cultures were incubated overnight at 28°C with shaking. The following day, the OD_600_ of the cultures was measured. The OD_600_ at the time of sampling did not differ between strains (p = 0.39 for BMM, p = 0.60 for stem homogenate; determined by Mann-Whitney tests). The OD_600_ values were used to calculate the sample volume needed to equal 1 mL of an OD_600_ 0.6 culture. The calculated volumes of each sample were centrifuged at 10,000 x g for 5 minutes at 4°C. The supernatant was removed and the cell pellet was immediately frozen in liquid nitrogen. Frozen samples were then shipped to the Mass Spectrometry and Metabolomics Core at Michigan State University (MSU).

At MSU, the extraction was performed as follows: 100 µL of extraction solvent (ACN/MeOH/H_2_O, 40:40:20, containing 50 nM c-di-GMP-F internal standard) was added to each tube containing a cell pellet. The tubes were vortexed and kept on ice for 30 minutes, followed by centrifugation for 10 min at 13,000 rpm. The supernatant was transferred to a new tube and the solvent was evaporated using a speed-vac. The sample was reconstituted in 100 µL of LC mobile phase (water containing 8 mM dimethylhexylamine and 2.8 mM acetic acid) containing 50 nM internal standard. Fully prepared samples were analyzed by LC-MS/MS on a Waters Xevo TQ-S triple quadrupole mass spectrometer. Calculated values are displayed as the concentration in the original 100 µL extract. Raw data are provided in Table S7.

### Lysozyme treatment

Strains prepared as described above were resuspended in 1 mL of water to an OD_600_ 0.1. Lysozyme from chicken egg white (Sigma-Aldrich L6876) was added to a final concentration of 12.2 µg/mL and suspensions were incubated statically at room temperature.

Control samples prepared in tandem did not have lysozyme added. After 30 min, samples were vigorously vortexed for 10 sec and then centrifuged at 8,000 x g for 5 min. The supernatant was removed and the cell pellets were resuspended in water before dilution plating to quantify surviving cell density.

### Statistical analysis

All statistics and area under the curve (AUC) calculations were performed using GraphPad Prism (version 10, GraphPad Software, San Diego, California USA). All datasets were first tested for normality and lognormality using the D’Agostino & Pearson test. If data fit a normal distribution, parametric statistical tests were performed. If data was non-normal but fit a lognormal distribution, statistical tests that compare the geometric means of each dataset were used. If neither test was passed, non-parametric tests were used. Specific statistical tests are indicated in each figure legend and p-values are reported in the graph or legend as appropriate.

## Acknowledgments

We gratefully acknowledge Shabda Gajbhiye and Mairead Morahan for their technical assistance. We further thank three anonymous reviewers for exceptionally constructive suggestions.

## Abbreviations

RSSC: Ralstonia solanacearum species complex
Rps: Ralstonia pseudosolanacearum
EPS: extracellular polysaccharide
ECM: extracellular matrix
c-di-GMP: bis-(3,5)-cyclic dimeric guanosine monophosphate

