## Supplemental Tables for "RprR is a plant-responsive regulator of exopolysaccharide production, biofilm formation, and virulence in *Ralstonia pseudosolanacearum*"

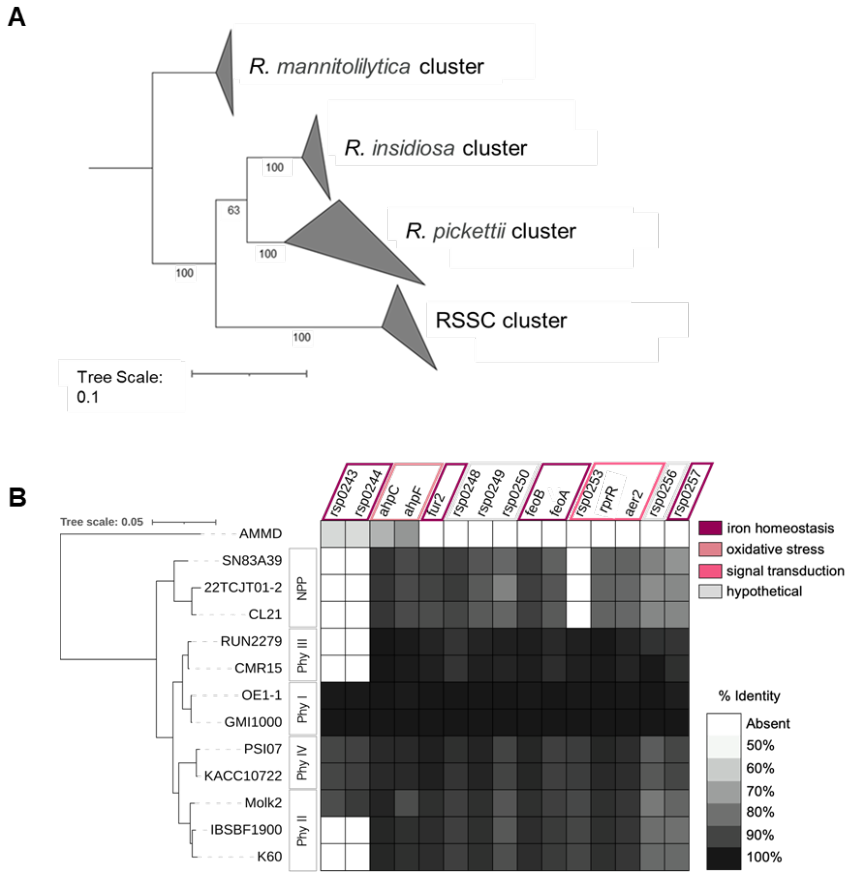


**Supplemental Figure S1: The *rprR* gene is conserved across the genus *Ralstonia*, along with a 12-gene syntenic cluster. A)** Maximum-likelihood gene tree constructed using the full-length *rprR* sequence from 106 *Ralstonia* genomes (See also Table S1 and supplementary methods). Individual branches were collapsed into 4 major taxonomic clusters. Bootstrap values are indicated on each node. Tree visualization was done in iTOL (1). **B)** Conservation of genes surrounding *rprR* in GMI1000. The protein blast function in KBase (2) was used to determine the conservation (≥50% protein sequence identity) of genes up- and downstream of *rprR* in GMI1000, indicated by degree of shading in the box representing each gene. The tree was edited in iTOL to overlay *rprR* conservation metadata and gene descriptions. Colored boxes outline genes based on the functional categories provided in the legend and mirror those shown in Figure 1A. **C)** Multiple-sequence alignment of a 34 amino acid region in the LOV domain. The comparison includes four representative RSSC strains (Plant Path) and three non-plant pathogenic (NPP) *Ralstonia* spp. The conserved 8 amino acid LOV motif is outlined by a black box. Yellow highlight indicates position 672 (GMI1000 as reference), which often encodes a highly conserved cysteine residue ($Cys_{672}$) where an oxidation-induced flavin-LOV adduct forms. The alignment was generated using Clustal Omega (accessed via EMBL-EBI web server (3, 4).

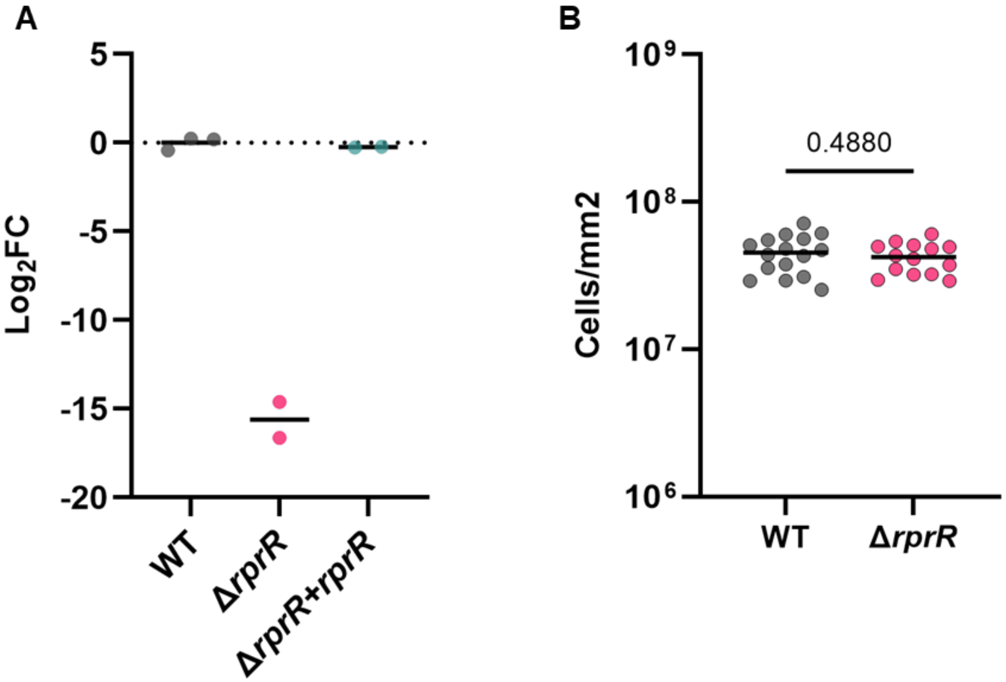


**Supplemental Figure S2: Expression of *rprR* is restored in the complemented mutant strain, and colony cell density on dilute rich media is not influenced by RprR. A)** Expression of *rprR in vitro*. Total RNA was extracted from cell pellets and qRT-PCR was used to quantify expression. Data is shown on a base-2 logarithmic scale relative to WT; the black dashed line indicates value '1' (mean WT expression). Each circle represents a biologically distinct culture (n=2-3 per strain). **B)** Colony cell density on dilute rich media. Each circle represents a single agar-grown colony. Lines represent the mean. The experiment was repeated three times with 2-6 colonies each (n=14-16 per treatment, p-value by Welch's T-test).

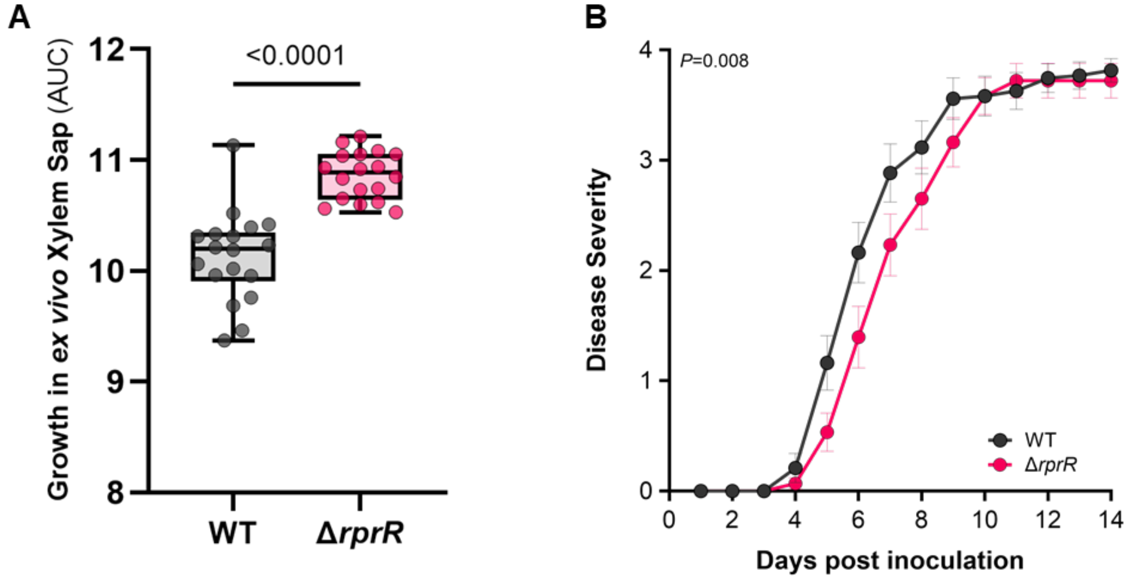


**Supplemental Figure S3: The Δ*rprR* mutant has a growth advantage in *ex vivo* xylem sap, but a virulence defect following a naturalistic soil-soak infection of a susceptible host.**

**A)** *Rps* growth in *ex vivo* xylem sap. Each circle represents the area under the curve (AUC) calculated for an individual microtiter well. Whiskers represent min to max values. Data shown represent 3 independent experiments, each with 6 technical replicates (n=18 per treatment, p-value by Welch's T-test). **B)** Wilt disease progress following soil-soak inoculation. Each symbol shows the mean disease index across three independent experiments, each containing 13-15 plants per treatment (n=43 per treatment, p-value by Repeated Measures two-way ANOVA). Error bars represent SEM.

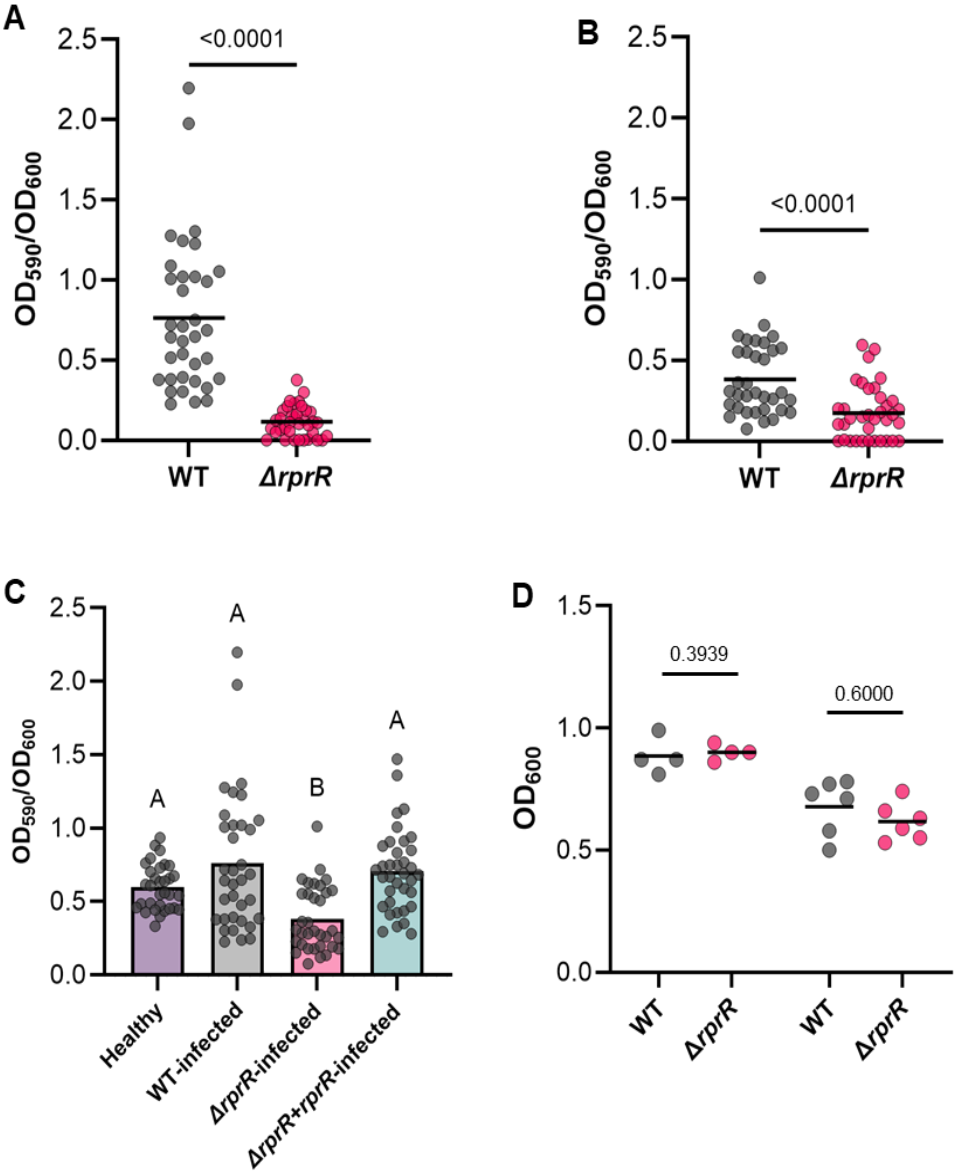


**Supplemental Figure S4: Infected tomato stem homogenate enables *Rps* biofilm formation *in vitro*, and samples used for c-di-GMP analysis were at similar cell densities.**

**A-B)** Biofilm formation in filter-sterilized stem homogenate from infected tomato plants. Tomato plants were infected three days prior to stem collection with either wild-type (A) or Δ*rprR* (B) cells. Each circle represents a single microtiter well. Lines represent the mean. Data reflect three independent experiments with 12 replicates each (n=36 per treatment). Outliers identified by a ROUT analysis (default parameters, implemented in GraphPad Prism) was removed prior to analysis and visualization. Data was analyzed using a Mann-Whitney test (A) and Welch’s T-test (B). **C)** Biofilm formation of *Rps* GMI1000 wild-type in filter-sterilized stem homogenate from healthy and infected tomato plants. Tomato plants were infected three days prior to stem collection with WT, Δ*rprR*, or Δ*rprR*+*rprR* cells, or left un-inoculated. Each circle represents a single microtiter well. Lines represent the mean. Data reflect three independent experiments with 12 replicates each (n=36 per treatment). Outliers identified by a ROUT analysis (default parameters, implemented in GraphPad Prism) were removed prior to analysis and visualization. Different letters indicate differences between groups (Lognormal Brown-Forsythe and Welch ANOVA). Data for healthy and WT-infected treatments is also shown in Figure 4C and S4A). **D)** Culture density of samples for c-di-GMP analysis. Each circle represents a single culture tube at the time of sample collection. Lines represent the mean. See Figure 6A for details on replicates. A Mann-Whitney test was performed for each comparison.

**Supplemental Methods**

***In silico* analyses of RprR.** *Ralstonia* genomes were obtained from the Joint Genome Institute's Integrated Microbial Genomes webserver (JGI IMG) in October 2023 (5). All genomes listed under the genus *Ralstonia* were manually curated to remove duplicate entries and assemblies with >60 scaffolds (final n = 111, Table S1). A "cassette search" of the genome set was performed using default parameters and the following hooks: pfam00672 (HAMP), pfam08447 (PAS_3), pfam13426 (PAS_9), pfam00990 (GGDEF), and pfam00563 (EAL). The hooks represent the JGI annotated pfam functions of the *Rps* GMI1000 *rprR* gene, excluding the CACHE domain, which we found was not consistently annotated within *rprR* across the genus. This functional search identified a single hit (hit = all hooks present within a 4kb genomic region) within 106 genomes. The amino acid sequence of the identified *rprR* genes (n = 106) were downloaded from JGI and subsequently uploaded to the CIPRES gateway for alignment using MUSCLE (default parameters, v3.7) (6, 7). The multiple-sequence alignment (MSA) was manually curated in Jalview (v2.11.2.7) (8) before building a maximum-likelihood tree in the CIPRES gateway (RAxML-HPC BlackBox, v8.2.12) (6, 9). The tree was uploaded to iTOL for visualization and formatting (1).

To visualize genomic context in Figure S1B, a phylogenetic tree was constructed on KBase (2) with 9 RSSC strains (representing all 4 phylotypes), 3 non-plant pathogenic (NPP) *Ralstonia* spp., and *Burkholderia ambifaria* AMMD as an outgroup. The phylogenetic analysis is based on 49 conserved genes (10). The protein blast function in KBase was used to determine the conservation (≥50% protein sequence identity) of genes up- and downstream of *rprR* in GMI1000.

**qRT-PCR.** To compare *rprR* gene expression across GMI1000 wild-type, Δ*rprR*, and Δ*rprR* + *rprR*, cultures were grown overnight in CPG to an $OD_{600}$ 0.2-0.6 (log phase) before pelleting via centrifugation. Total RNA was isolated from the cell pellets using a Direct-zol RNA extraction kit (Zymo Research). RNA was reverse transcribed into cDNA with the SuperScript VILO cDNA Synthesis Kit (Invitrogen). A targeted qRT-PCR measured expression of *rprR* and two normalization genes (*rplM* and *serC*) using primers listed in Table S5. The ΔΔCt method was used to calculate relative gene expression in Δ*rprR* and Δ*rprR* + *rprR* compared to wild-type.

**Agar-grown colony cell density.** Strains were prepared as described in the main methods and resuspended to a concentration of approximately 50-100 CFUs/mL in water. A 100 uL volume of each suspension was spread on an agar plate containing a modified dilute rich media (2 g/L peptone, 0.2 g/L casamino acids, 1.8 g/L glucose, 0.2 g/L yeast extract and incubated in the dark at 28°C. After 4 days, the plates were imaged and two orthogonal measurements of each colony diameter were taken and averaged. This measurement was used to calculate the area of the colony ($A = \pi r^2$). The measured colonies were then cored from the plate using the top-end of a 1,000 uL pipet tip, homogenized in water, and serially diluted.

**Growth in *ex vivo* xylem sap.** Xylem sap was collected as described in the main methods. Cultures were prepared as described in the main methods and resuspended to a final $OD_{600}$ of 0.01 in thawed xylem sap. Cell suspensions were aliquoted into flat-bottomed 96-well plates (200 µL/well, Corning Costar Ref# 3370) and incubated at 28°C with slow continuous shaking in a Biotek plate reader. The $OD_{600}$ was measured every 30 minutes for 24 hours.
